# Rectus femoris hyperreflexia predicts knee flexion angle in Stiff-Knee gait after stroke

**DOI:** 10.1101/699108

**Authors:** Tunc Akbas, Kyoungsoon Kim, Kathleen Doyle, Kathleen Manella, Robert Lee, Patrick Spicer, Maria Knikou, James Sulzer

**Author notes:** Corresponding Author: James Sulzer, PhD, 204 E. Dean Keeton St, Austin, TX 78712.

## Abstract

Stiff-knee gait (SKG) after stroke is often accompanied by decreased knee flexion angle during the swing phase. The decreased knee flexion has been hypothesized to originate from excessive quadriceps activation. However, it is unclear whether this activation is due to poor timing or hyperreflexia, both common post-stroke impairments. The goal of this study was to investigate the relation between quadriceps hyperreflexia in post-stroke SKG with knee flexion angle during walking. The rectus femoris (RF) H-reflex was recorded in eleven participants with post-stroke SKG and ten healthy controls during standing and walking during toe-off. In order to separate the effects of poorly timed quadriceps muscle activation from hyperreflexia, healthy individuals voluntarily increased quadriceps activity using RF electromyographic (EMG) biofeedback during standing and pre-swing upon H-reflex stimulation. We observed a negative correlation (R = −0.92, p=0.001) between knee flexion angle and RF H-reflexes in post-stroke SKG. In contrast, H-reflex amplitude in healthy individuals in presence (R = 0.47, p = 0.23) or absence (R = −0.17, p = 0.46) of increased RF activity had no correlation with knee flexion angle. The RF H-reflex amplitude differed between standing and walking in healthy individuals, including when RF activity was increased voluntarily (*d* = 2.86, p = 0.007), but was not observed post-stroke (*d* =0.73, p = 0.296). Thus, RF reflex modulation is impaired in post-stroke SKG. Further, RF hyperreflexia, as opposed to overactivity, may play a role in knee flexion kinematics in post-stroke SKG. Interventions targeting self-regulated quadriceps hyperreflexia may be effective in promoting better neural control of the knee joint and thus better quality of walking post-stroke.

## Introduction

Stiff-Knee gait (SKG) is one of the most common gait disabilities following stroke. SKG is defined as reduced knee flexion (Perry & Burnfield, 1992) and decreased knee flexion velocity (Goldberg et al., 2003) during the swing phase of the gait. Those with SKG have reduced walking function, joint pain (Hsu et al., 2003) and energy inefficiency (Doke et al., 2005; Royer & Martin, 2005; Shorter et al., 2017). Post-stroke SKG has been primarily attributed to quadriceps overactivity (Kerrigan et al., 1991) and poorly timed muscle co-activation profiles (Riley & Kerrigan, 1998) based on early studies measuring muscle activation. Later, more sophisticated biomechanical analyses focused on knee flexion velocity (Anderson et al., 2004; Goldberg et al., 2006; Goldberg et al., 2003), inappropriate ankle plantarflexion torque (Lamontagne et al., 2000), increased quadriceps forces (Goldberg et al., 2004), knee extension moment prior to swing phase (Burpee & Lewek, 2015; Kerrigan et al., 1997) and simulated joint moments (Piazza & Delp, 1996) and muscle forces (Piazza, 2006). However, the specific neuromuscular mechanisms causing SKG after stroke remain unclear.

While quadriceps overactivity is the most widely believed cause of SKG (Kerrigan et al., 1991; Reinbolt et al., 2008; Sutherland et al., 1990), it is unclear why such overactivity would occur. Increased quadriceps activation could be achieved voluntarily to improve stance leg stability but fail to deactivate in time for pre-swing phase (Higginson et al., 2006; Reinbolt et al., 2008). Quadriceps activation could also occur through hyperreflexia or tone occurring during pre-swing quadriceps stretch caused by knee flexion (Akbas et al., 2019; Faist et al., 1999). Researchers have used Botox injections to deactivate the femoral nerve, showing modest improvements in knee flexion (Robertson et al., 2009; Roche et al., 2015; Stoquart et al., 2008). These results suggest that overactivity plays a role in SKG but cannot ascertain its source. Understanding the specific mechanisms of SKG could help develop targeted interventions.

Evidence from our earlier studies indicates that hyperreflexia could be a main factor in poststroke SKG. Specifically, we used a custom-designed robotic knee flexion actuator (Sulzer et al., 2009) to perturb knee flexion in nine individuals with post-stroke SKG and five healthy controls (Sulzer et al., 2010) during pre-swing. While knee flexion angle increased smoothly in healthy controls with the perturbation, we observed a sharp knee extension following initial increased knee flexion in post-stroke SKG. Integrated EMG extracted within 100 ms following the perturbation revealed consistently increased RF EMG within participants (Akbas et al., 2017). Further analysis with musculoskeletal modeling and simulation estimated that within a period reflecting reflex latency, increased RF activity followed maximum RF fiber stretch velocity (Akbas et al., 2019). Taken together, these data suggest that RF hyperreflexia is a consistent factor in those with post-stroke SKG. However, it is unclear to what degree RF hyperreflexia is responsible for reduced swing-phase knee flexion.

Hyperreflexia in the muscles of the knee and ankle are well-known after stroke (Given et al., 1995; Marque et al., 2001; Maupas et al., 2004; Zehr et al., 1998). For example, Faist et al., (1999) found hypersensitive quadriceps tendon jerk reflexes during post-stroke gait. Elevated involuntary responses in quadriceps has been observed by implementing mechanical perturbations in hip flexion (Lewek et al., 2007; Lewek et al., 2006) and hip abduction (Finley et al., 2008). These altered involuntary responses could play a role in diminished knee flexion during the swing phase of post-stroke SKG. However, mechanical perturbations introduce additional intermediate factors mediating monosynaptic reflex responses. For instance, knee flexion perturbations during gait could induce kinetic interaction and torque couplings between different joints (e.g. knee and ankle) and result in a time delay between the initiation of reflex and perturbation (Mrachacz-Kersting, Lavoie, Andersen, & Sinkjaer, 2004).

In contrast, a monosynaptic reflex can be directly elicited with a H-reflex by peripheral nerve stimulation, bypassing gamma motoneurons (Pierrot-Deseilligny & Burke, 2005). The stimulation can measure the monosynaptic hyperexcitability in Group Ia afferent pathways (Knikou, 2008). Thus, as a probe, electrically evoking H-reflex is more reliable and consistent in identifying the altered neuromuscular pathways, especially for identifying the reflex sensitivity and modulation in neural control of movement (Misiaszek, 2003). The modulation of H-reflexes in knee extensors and ankle extensors have been well documented (Dietz et al., 1990) and associated to joint angles of knee and ankle during gait (Larsen et al., 2006; Sinkjær et al., 1996). Enhanced intersegmental facilitation of soleus muscle from hyperactive quadriceps afferents has been revealed after stroke (Dyer et al., 2009; Dyer et al., 2011) and related to altered activation timings of knee and ankle extensors during gait following stroke (Dyer et al., 2014). However, altered reflex excitability in the targeted muscle is highly context-dependent. For example, increased excitation of knee extensors was induced by peroneal nerve stimulation during walking (Achache et al., 2010), whereas the same stimulation suppressed quadriceps excitability during knee flexion while pedaling (Fuchs et al., 2011) following stroke. It is unknown how quadriceps reflex excitability is modulated during gait in those with post-stroke SKG.

The objective of this study was to characterize the role of hyperreflexia on knee flexion in post-stroke SKG. Specifically, our goals were to determine how monosynaptic quadriceps reflex responses were modulated in standing and walking, and whether the magnitude of such reflex responses could be related to the degree of knee flexion after stroke. We evoked RF H-reflex in both post-stroke individuals and healthy controls. In order to delineate reflex modulation due to hyperreflexia from voluntary overactivity, we compared responses in individuals who were poststroke with healthy controls asked to increase RF muscle activity voluntarily using EMG biofeedback. We expected elevated RF H-reflex sensitivity during both standing and walking (i.e. during toe-off) in people with post-stroke SKG compared to healthy controls. We also predicted that the level of hyperexcitability of the RF H-reflex would be associated with diminished knee flexion during swing phase in SKG. Characterization of the role of hyperreflexia in SKG and other neuromuscular disorders could help to design more targeted treatments for hyperreflexia such as operant H-reflex conditioning (Thompson et al., 2009).

## Methods

### Data Collection

Eleven hemiparetic individuals with post-stroke SKG (9 M/ 2 F, 53±9 years mean±SD, Table 1) able to walk without assistance for fifteen minutes, at least six months following injury with exclusion of multiple strokes, cerebellar damage, lower limb musculoskeletal injury written informed consent according to the guidelines approved by the local Institutional Review Board to participate in the experiment. Post-stroke participants whose knee range-of-motion (ROM) was least 20° less on the affected side compared to the unaffected side qualified as SKG. Participants also had to be able to walk 20 minutes at minimum of 0.2 m/s on a treadmill. Ten healthy controls (6 M/ 4 F, 35±13 years mean±SD) with no musculoskeletal injuries also completed the protocol. All subjects walked on an instrumented split-belt force treadmill (Bertec, Columbus, OH), which recorded ground reaction forces. Lower limb kinematic data were collected using inertial motion capture (IMU, Xsens, Enschede, Netherlands). Surface EMG (Bortec, Calgary, AL) were collected with electrodes placed on the muscle belly of adductor longus (AL), GMed, tensor fascae latae (TFL), gluteus maximus (GMax), medial amstrings (MH), RF and vastus medialis (VM). IMU data was collected at 60 Hz, whereas the other data were collected at 1 kHz. In one post-stroke individual, we were unable to record motion data due to equipment failure, and thus for the participant’s data was not included only in subsequent kinematic analysis.

**Table 1.** Data for 11 participants with post stroke Stiff-knee gait

| Subject no | Age (yrs) | G | Side | W (kg) | Post (yrs) | Med. | Botox | AFO/ Cane | Walking Speed (m/s) | Peak Knee flexion (degree) |
| --- | --- | --- | --- | --- | --- | --- | --- | --- | --- | --- |
| 1 | 41 | M | L | 100 | 3 | - | No | - | 0.5 | 26.3 |
| 2 | 70 | M | R | 86 | 4 | Baclofen | No | AFO | 0.35 | 6.9 |
| 3 | 63 | M | R | 81 | 1.5 | - | Yes | Cane | 0.5 | 19.5 |
| 4 | 55 | M | L | 110 | 8 | Baclofen | Yes | Cane | - | - |
| 5 | 45 | M | L | 76 | 2.5 | - | No | AFO | 0.35 | 11.8 |
| 6 | 59 | M | R | 83 | 1 | - | Yes | - | 0.3 | 9.9 |
| 7 | 53 | M | R | 65 | 3 | - | Yes | - | 0.3 | 12.6 |
| 8 | 53 | M | R | 77 | 1 | - | Yes | - | 0.4 | 15.6 |
| 9 | 41 | F | R | 64 | 7 | - | No | - | 0.5 | 27.7 |
| 10 | 61 | F | L | 63 | 1 | - | No | Cane | 0.22 | 9.5 |
| 11 | 47 | M | L | 89 | 2 | - | Yes | - | - | - |
| Mean | 53.4 |  |  | 81.3 | 3.1 |  |  |  | 0.38 | 15.5 |
| SD | 9.4 |  |  | 14.9 | 2.4 |  |  |  | 0.10 | 7.5 |

\*All the AFOs wore by the participants were hinged versions. "Walking Speed" refers to each participants maximum comfortable walking speed on treadmill. Subject 4 was only able to walk over ground and walking speed and peak knee flexion angle measures were not collected
G indicates gender; W, weight; Post, years post-stroke; Meds, taking Baclofen; AFO/Cane, use of an ankle-foot orthosis or cane; M, male; F, female.

The experimental protocol began with recruitment of the RF H-reflex during standing (Figure 1 a-b), and followed by repetitive stimulations, seven seconds apart from each other to avoid post-activation depression (Hultborn et al., 1996; Pierrot-Deseilligny & Burke, 2005) of the femoral nerve. The stimulations were applied at the current associated with H_max_ during standing in four different conditions: standing posture (*Stand*), walking at toe-off (*Walk*), standing with increased RF EMG activity (*Stand*↑) and walking at toe-off with increased EMG activity (*Walk*↑). These conditions are illustrated in Figure 2. For each participant, repetitive stimulations were implemented 25-30 times during each condition with the intensity corresponding to maximum RF H-reflex (H_max_) obtained from the recruitment. The repetitive stimulations were applied in pseudo-random steps, seven to nine seconds apart from each other, during standing conditions and during toe-off, matching the timing of the knee flexion perturbation in our previous study (Sulzer et al., 2010). The phases were detected automatically in real-time using the vertical ground reaction force measures of the affected side from treadmill. Those with post-stroke SKG were allowed to grip the supporting handrails of the treadmill to maintain balance. A harness was used to prevent falls, and the participants walked at the lesser of the maximum achievable walking speed or 0.5 m/s for 15 minutes for the walking conditions. In participants with SKG, the *Stand* and *Walk* conditions were in randomized order, whereas the *Stand↑* condition was optional due to fatigue and the *Walk*↑ condition was not included due to the difficulty of the task. Two of the stroke participants were able to complete the *Stand*↑ condition. For healthy controls, the walking speed was fixed at 0.5 m/s for the walking conditions and all the conditions were randomized. The corresponding H_max_ measures were normalized (H_norm_) with respect to maximum motor responses (M_max_) for comparisons between conditions and groups.

**Figure 1.**
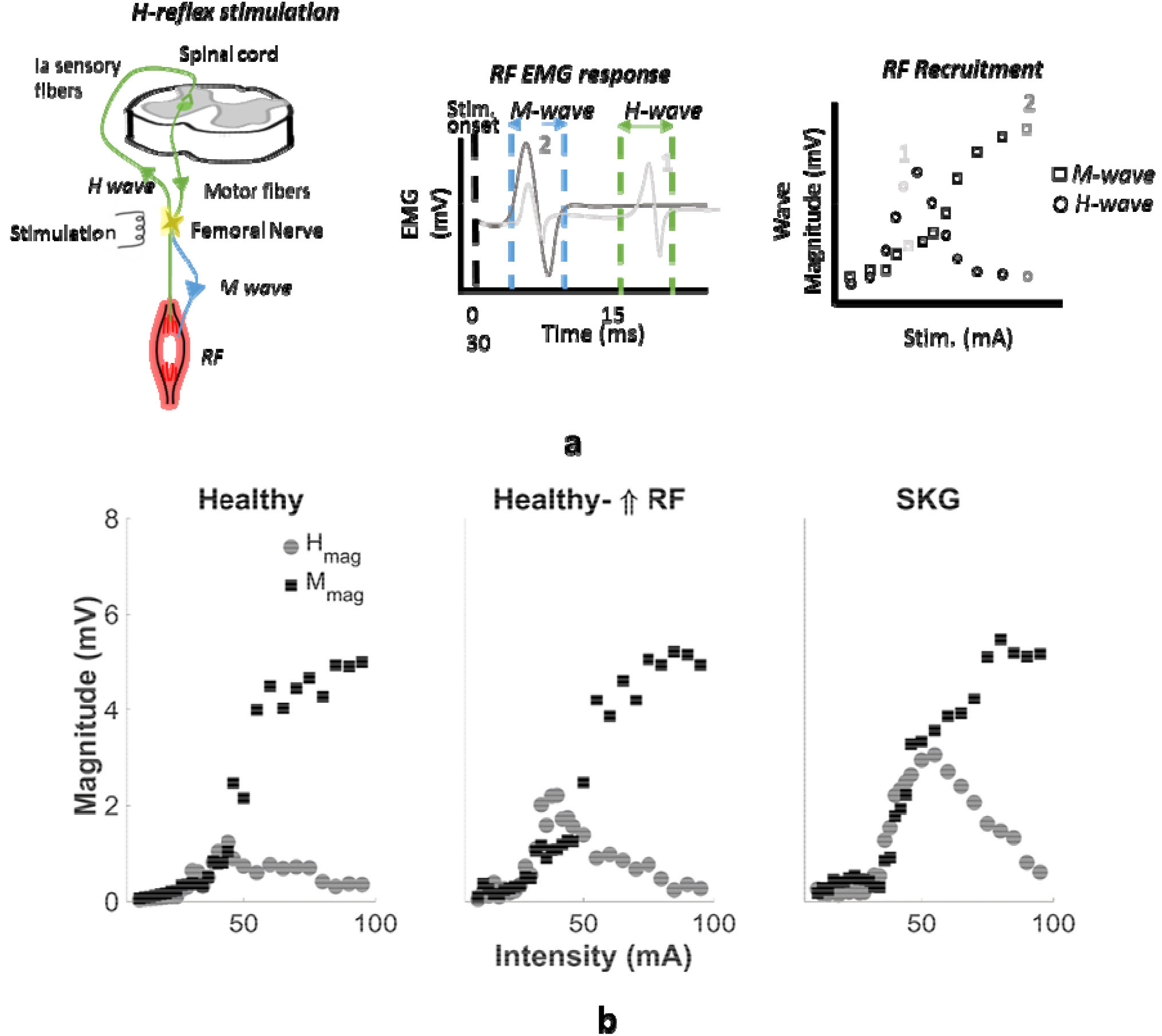
RF H-reflex recruitment process (top) and representative H-reflex recruitment curves for each group. H-reflex recruitment process starts by applying gradually increasing electrical stimulation on femoral nerve (top left) in 2 mA increments between 10-46 mA and 5 mA between 50-90 mA with two stimulation trials for each interval in standing posture. Based on the response time of EMG responses, the optimal H-reflex response (H-wave), and maximum motor response (M-wave) (top middle) and corresponding recruitment curve (top right) was obtained. Compared to healthy baseline representative (bottom right) the recruitment curves indicate increased RF H-wave in healthy representative with increased voluntary RF contraction (bottom middle) and in participant with post-stroke SKG (bottom right). No considerable change was observed between maximum M-waves (M_max_) between conditions in healthy representative

### RF H-reflex recruitment

We applied stimulation to the femoral nerve using a stimulator (Digitimer DS7A, Hertfordshire, UK). The optimal nerve location was detected by applying stimulation to a grid of points covering femoral triangle starting from the previously referred anatomical landmark (Zehr, 2002) started with a fixed intensity (30 mA) matching the average maximal H-reflex response from the literature (Doguet & Jubeau, 2014; Larsen & Voigt, 2006). After the femoral nerve location was determined, the cathode electrode was placed on the femoral nerve and anode electrode was placed on the soft tissue on back upper thigh of the affected/matched side. Afterwards, the RF H-reflex recruitment curves were obtained in a standing posture by applying gradually increasing electrical stimulation trials on the femoral nerve (in 2 mA increments between 10-46 mA and 5 mA between 50-90 mA with two stimulation trials for each interval). The representative recruitment curves for a healthy control and a participant with post-stroke SKG are illustrated in Figure 1b.

Recruitment of the quadriceps H-reflex curves were challenging due to the occasional overlap between the M- and H-waves. This issue was addressed previously with a post-hoc analysis by subtracting M_max_ from the trials with H-reflex response (Larsen et al., 2006; Larsen & Voigt, 2006). However, due to the custom-defined selection of the descending portion of M_max_ for the subtraction in subjects and the time and magnitude difference between the EMG responses, this post-hoc analysis is vulnerable to the variability between subtracted signals. As an alternative, we introduced an approach which selects the best decaying exponential fit starting from the peak M-wave by least root-mean square error and subtracted the fitted signal from the raw H-wave (H_raw_) signal to obtain corrected H-wave (H_corr_) (Figure 3). Afterwards we stored the M_max_ and H-wave with the highest magnitude at the ascending portion of the recruitment curve (H_max_) to guarantee monosynaptic reflex response from Group Ia afferents (Knikou, 2008, Figure 1a). The corresponding intensities for H_max_ responses were stored and used for repetitive trials in conditions. One post-stroke participant was excluded from the study due to a recent Botox injection and subsequent inability to evoke an RF H-reflex.

### RF EMG feedback

Volitional activity of the RF was increased for *Stand*↑ and *Walk*↑ conditions using RF EMG signal as a feedback for controls (Figure 2). For standing conditions, the rectified RF EMG was displayed with an onset cue provided 300 ms prior to stimulation. The participants were instructed to increase their RF EMG signal above the displayed continuous threshold value, set to be 20% of RF maximum voluntary contraction (MVC) obtained by isometric contractions during standing posture. The trials were considered “successful” if the increased rectified RF EMG exceeded the threshold and the condition continued until 25 successful trials were obtained (Figure 2-*Stand*↑). The standing feedback condition remained optional for participants with post-stroke SKG and always implemented at the end of the experiment and only for the standing conditions. For the walking feedback conditions of the controls, the onset cue was set to 20 ms prior to toe-off and displayed for every gait cycle with the same threshold value in standing. The repetitive stimulations were applied during pseudo-random steps, at least 5 seconds apart from each other, and continued until 25 successful trials were obtained (Figure 2-*Walk*↑). Since the subjects were required to adjust their walking in order to achieve increased voluntary RF contraction, an adaptation period without nerve stimulations was implemented until the participants were able to successfully complete ten consecutive steps with voluntary RF contraction. One healthy participant was removed from analysis due to inability to achieve the task.

**Figure 2:**
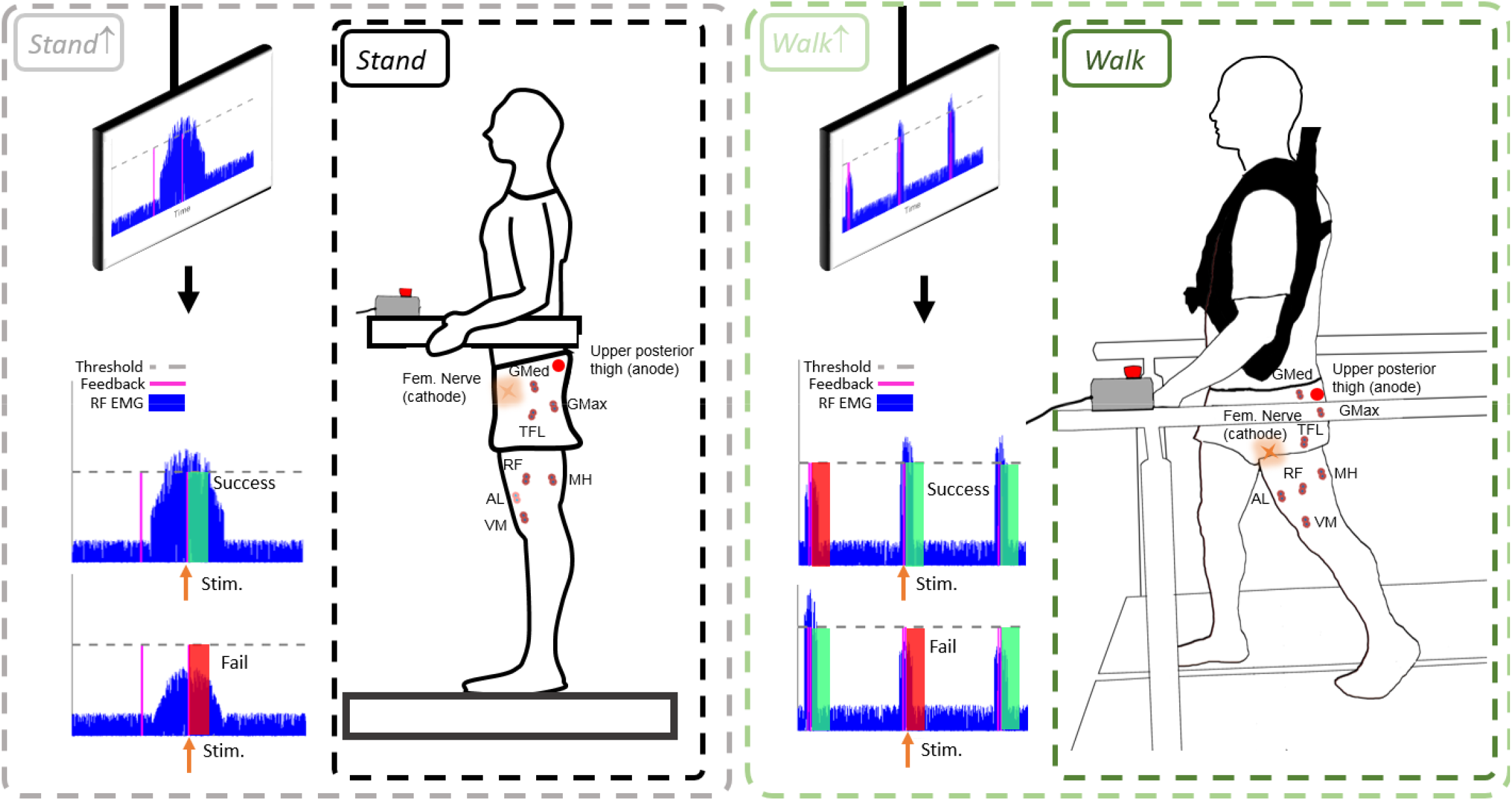
Experimental protocol and RF EMG feedback paradigm. The dashed square boxes indicate the four conditions in experimental protocol namely *Stand*, during walking at toe-off *Walk*, during standing with increased RF EMG activity *Stand*↑ and during walking at toe-off *Walk*↑. For each condition 25-30 repetitive stimulations corresponding to maximum H-reflex response were applied. All conditions were implemented in randomized order. For *Stand*↑ and *Walk*↑, the participants were instructed to increase their RF EMG signal above the threshold value (dashed gray line), 20% of RF maximum voluntary contraction (MVC) following a que signal in real-time (purple vertical line) with 300 ms and 20 ms offset time prior to stimulation respectively. For condition *Walk*↑ the que has been displayed in every gait cycle to account for dynamic adaptation during walking. The trials were considered “successful” if the increased rectified RF EMG exceeds the threshold during stimulation (top figures in *Stand*↑ and *Walk*↑) and considered “fail” otherwise. (Bottom figures in *Stand*↑ and *Walk*↑)

**Figure 3:**
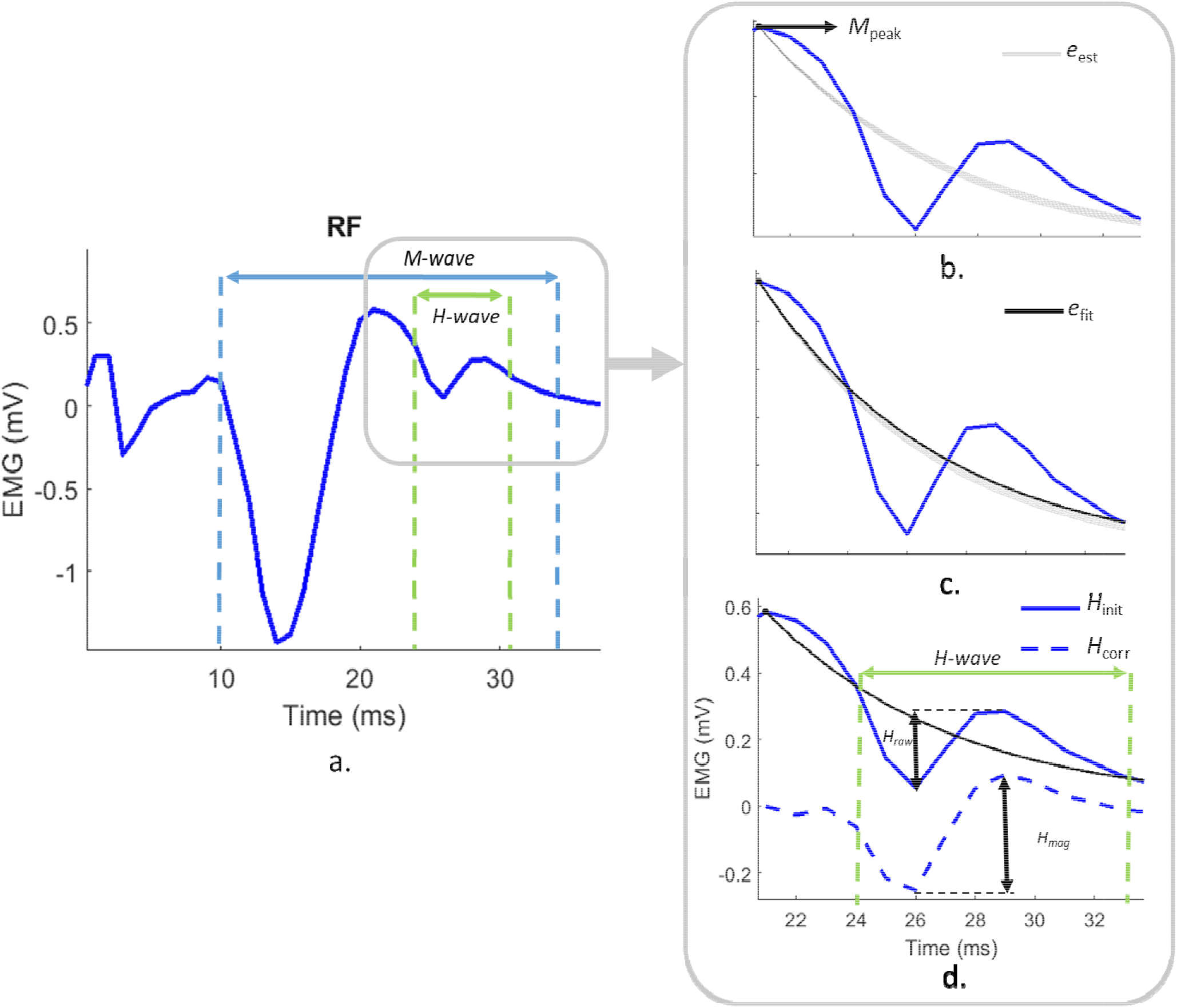
Post-hoc analysis for RF H-reflex magnitude extraction. The post-hoc analysis applied on the corresponding RF EMG responses where H-wave overlapped with descending portion of M-wave (a). First, the exponential decaying fit lines (e_est_) with least root mean squared error (RMS) starting from the peak M-wave and ending around the vicinity of convergence point were determined (b). From those, the exponential fit with the least average RMS was selected (c) and subtracted from the initial H-wave segment (H_ini_t) to obtain corrected H-wave segment (H_corr_) (d) to obtain the isolated H-reflex response (H_mag_) from raw H-reflex response (H_raw_).

### Statistical Analysis

We extracted the peak-to-peak H-reflex amplitude normalized to M_max_ and knee flexion ROM during swing phase as the main outcome measures. We used linear-mixed effect models with Tukey post-hoc t-tests (α < 0.05) to examine differences between conditions. Conditions within healthy individuals were paired accordingly. Correlation coefficients between knee flexion ROM and H-reflex amplitude were obtained using Pearson correlation. All statistics were performed using the R statistical package.

## Results

We observed a strong relationship between peak knee flexion during swing phase and RF H-reflex measures in post-stroke SKG (R = −0.92, p < 0.001). No correlation was observed both for healthy *Walk* (R = −0.17, p = 0.46) or healthy *Walk*↑ (R = 0.47, p = 0.23) conditions. However, it should be noted that one healthy individual was defined as an outlier and removed due to RF activity 2.5 SD above the mean in *Walk*↑, resulting in N=8 healthy individuals. Results are illustrated in Figure 4. The relation in SKG was different compared to healthy *Walk* (t = 5.81, p = 0.028) and healthy *Walk*↑ (t = 9.33, p = 0.011). There was no significant difference observed between healthy walking conditions (t = 0.75, p = 0.491).

**Figure 4:**
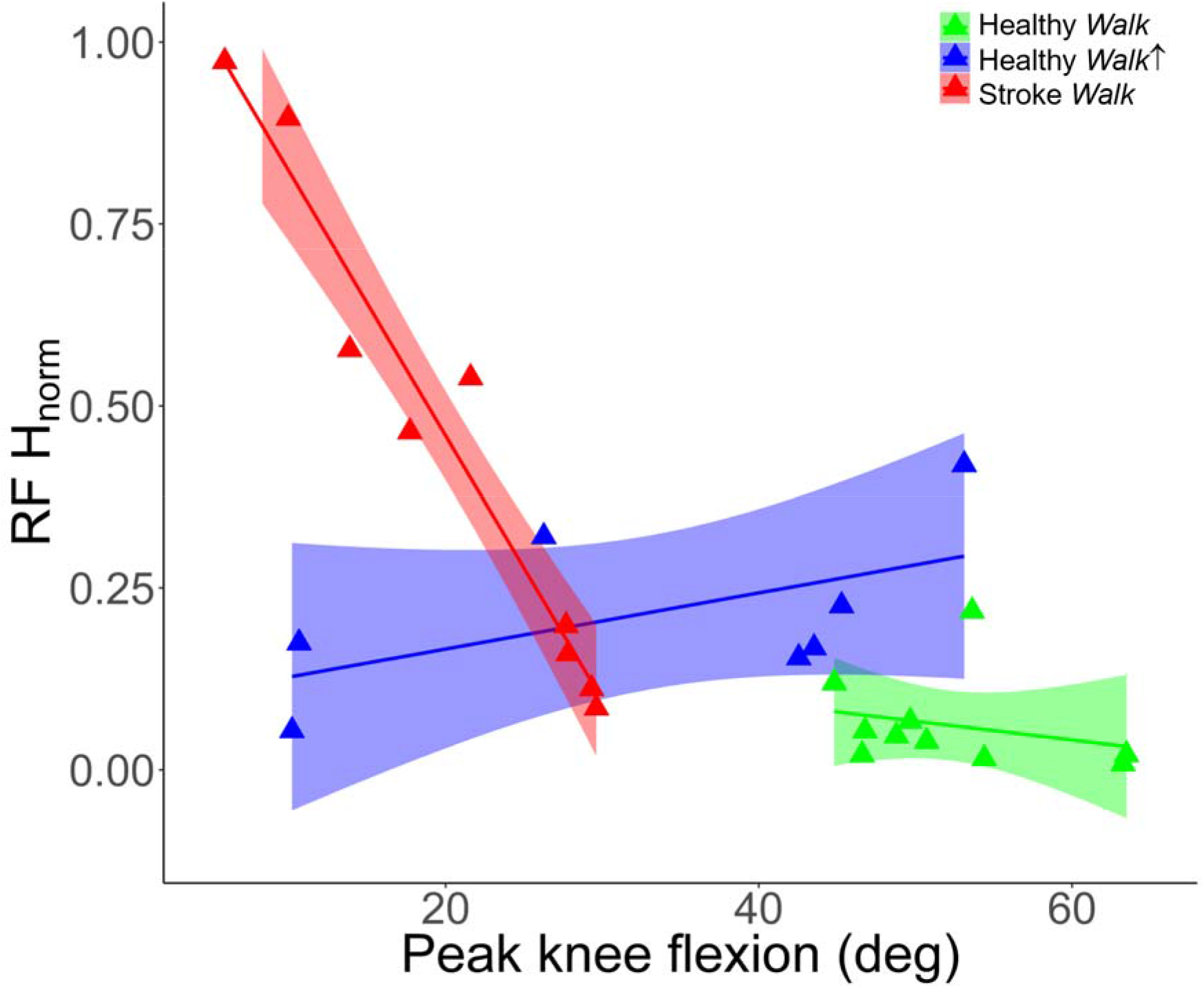
Diminished knee flexion is correlated with increase RF H-reflex in post stroke SKG. Peak knee flexion angles and RF H-reflex magnitudes indicate significant correlation in participants with post-stroke SKG (red triangles, *R* = −0.92) with decreased knee flexion during swing phase whereas no significant correlation was observed in healthy *Walk* (green triangles,, *R* = −0.17) and healthy *Walk*↑ (blue triangles,, *R* = 0.47). The interaction between RF H-reflex and knee flexion in SKG was significant compared to healthy *Walk* (*p* < 0.028) and healthy *Walk*↑ (*p* = 0.011). There was no significant difference in RF H-reflex and knee flexion between healthy groups (*p* = 0.75). The lines and shaded areas indicate the linear regression fits and 95% confidence interval respectively for the corresponding groups.

During standing, RF H-reflex measures in post-stroke SKG (N=10) were not significantly different than healthy controls in *Stand*↑ (t = 1.93, p = 0.071). Both the SKG group and healthy *Stand*↑ RF H-reflex measures were greater than healthy *Stand* (t = 3.51, p = 0.003, t = 13.82, p < 0.001, respectively, Figure 5). For walking conditions, RF H-reflex was higher in post-stroke SKG compared to healthy *Walk* (t = 3.28, p =0.004) and *Walk*↑ (t= 2.17, p = 0.044, Figure 5, right). RF H-reflex was also higher in healthy *Walk↑* compared to healthy *Walk* (t= 15.35, p < 0.001).

**Figure 5:**
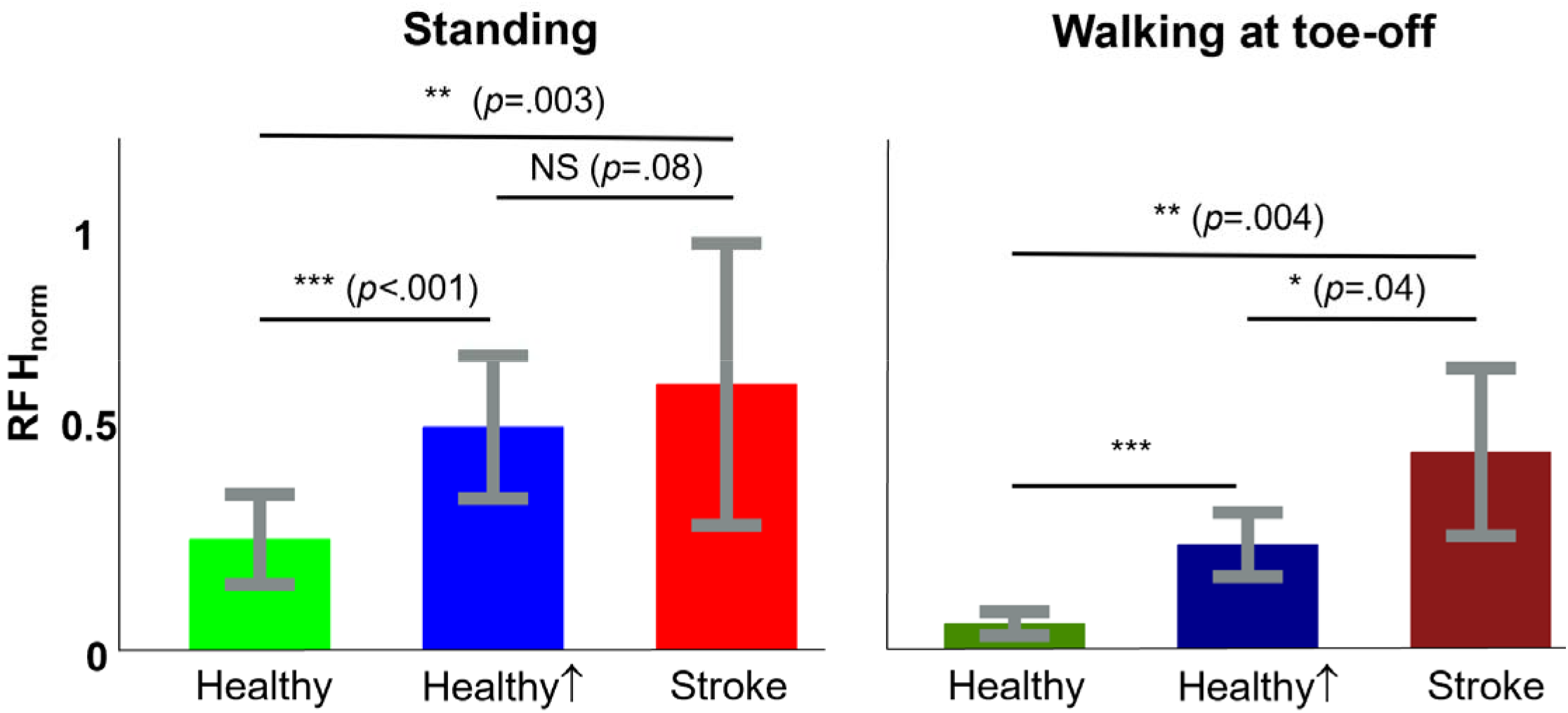
Elevated RF H-reflex in post-stroke SKG is higher than increased voluntary RF contraction. During standing, RF H-reflex response were significantly increased in post-stroke SKG (p = 0.003) and healthy groups with increased voluntary RF contraction (p < 0.001) compared to healthy baseline, whereas no significant difference was found between post-stroke SKG and increased RF activity (p = 0.08). During walking however, RF H-reflex response was increased in post-stroke SKG compared to both healthy baseline (p = 0.004) and with increased voluntary contraction (p = 0.04).

We found increased RF H-reflex measures in *Stand* compared to *Walk* (t= 3.23, p = 0.014), and *Stand*↑ compared to *Walk*↑ (t= 3.78, p = 0.007) in healthy controls, whereas no change was observed between *Stand* and *Walk* in participants with post-stroke SKG (t= 1.10, p = 0.296). Results are shown in Figure 6.

**Figure 6:**
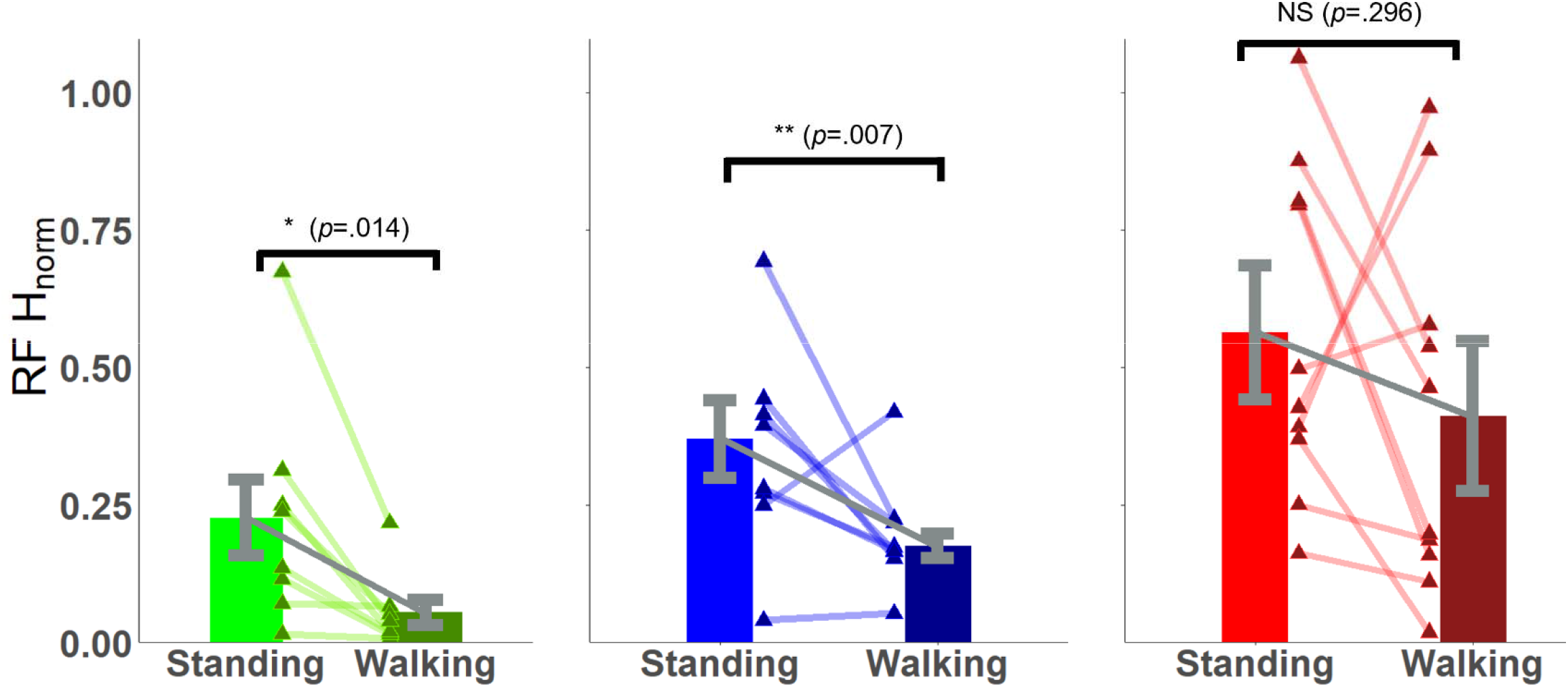
RF H-reflex modulation during walking is absent in post-stroke SKG. RF H-reflex magnitudes were significantly decreased in healthy controls between *Stand* and *Walk* (p = 0.014), and *Stand*↑ and *Walk*↑ (p = 0.007) conditions. There was no significant difference in participants with post-stroke SKG (p = 0.296).

## Discussion

Our goal was to determine the influence of RF hyperreflexia on reduced knee flexion during post-stroke SKG. We distinguished the effect of increased voluntary quadriceps contraction on RF H-reflex modulation by increasing EMG activity in healthy controls during standing and walking. Our results revealed a strong correlation between elevated RF H-reflex at toe-off and reduced peak knee flexion during swing phase in SKG. We observed no such relation in healthy individuals walking normally or with increased RF activation. H-reflex was modulated in in healthy controls between walking and standing, even with increased RF activation, whereas no such modulation was found in post-stroke SKG. The RF H-reflex amplitude was higher during toe-off in post-stroke SKG than healthy individuals with or without increased RF activity. Together, these findings provide new evidence that quadriceps hyperreflexia contributes to gait impairment in post-stroke SKG.

Overactivity of quadriceps, particularly RF, has been believed to be the primary cause of post-stroke SKG (Kerrigan et al., 1997; Riley & Kerrigan, 1998; Lewek et al., 2007). The consequences of such activation have been attributed to reduced knee flexion velocity in swing phase (Campanini et al., 2013; Goldberg et al., 2004; Goldberg et al., 2003), as well as excessive knee extension moment during pre-swing phase (Goldberg et al., 2006) in those with SKG. However, it is unclear whether such excessive quadriceps activity is due to reflexive mechanisms. We elucidated a strong correlation between quadriceps reflex excitability and reduced knee flexion during swing in those with post-stroke SKG. Further, this correlation was independent of reflex excitability during simulated voluntary “overactivity” of the quadriceps in healthy individuals, indicating that quadriceps reflex excitability may be a mechanism of SKG. This finding concurs with previous studies correlating increased knee extension velocity with SKG (Damiano et al., 2006). Thus, our findings suggest that it is unlikely that poorly timed, graded or coordinated quadriceps activation is responsible for reduced knee flexion, but instead a hyperactive quadriceps monosynaptic reflex is the more likely culprit.

We found context-dependent RF H-reflex modulation in healthy individuals between standing and walking but no change in RF H-reflex responses between standing and walking in SKG. This finding was in agreement with reduced quadriceps tendon jerk reflex modulation between standing and walking in post-stroke gait (Faist et al., 1999). The conserved elevated RF H-reflex magnitudes in SKG reveals the translation of previously reported quadriceps H-reflex responses during rest (Marque et al., 2001; Nielsen et al., 1995) to locomotion which is similar to the translation of elevated soleus H-reflex responses during standing (Hwang et al., 2004) and cycling (Fuchs et al., 2011; Schindler-Ivens et al., 2008). In healthy locomotion, quadriceps and soleus H-reflex modulation are phase-and task-dependent (Capaday & Stein, 1986; Larsen & Voigt, 2006; Lavoie et al., 1997) but not influenced by restricted knee movement during gait (Schneider et al., 2000). Here we additionally found that increased voluntary activation of the RF also did not influence such reflex modulation in healthy individuals, suggesting that overactive quadriceps activation was not responsible for the lack of modulation observed in SKG. Hence, the absence of H-reflex modulation between standing and walking in SKG could indicate an alteration in central neural control, likely arising from subcortical structures such as the reticulospinal or vestibulospinal tracts (Li & Francisco, 2015).

While the correlation between elevated H-reflex and decrease in knee flexion introduces a specific neurophysiological factor to evaluate reduced knee flexion in SKG, there could be other causes of reduced knee flexion. This study does not exclude the influence of heterogeneous abnormal mechanisms such as altered coordination between unaffected and affected knee extensors (Kautz & Patten, 2005), altered heteronymous reflex pathways influencing knee extensors and ankle flexors (Dyer et al., 2009), and simultaneous involuntary knee extensor and hip adductor activity (Finley et al., 2008), all of which could influence knee movement in SKG. The evaluation of these mechanisms introduces additional practical challenges such as concurrent nerve stimulation or customized perturbations for specific joint movements during walking. More importantly, the influence of homonymous pathways on individual joint movements during gait is necessary to establish prior to more complex coordinated joint movements governed by heterogeneous neural mechanisms. The revealed relation between elevated monosynaptic quadriceps stretch reflex responses and knee flexion is a crucial step towards establishing the influence of homonymous pathways in SKG.

This study also has practical limitations preventing the accurate imitation of over-active quadriceps in SKG on healthy controls. The increased RF activity in standing and walking was limited to voluntary activation generated by healthy motor control and did not accurately represent the influence of altered intrinsic muscle properties (Horstman et al., 2008) and the altered quadriceps activity originated from reduced muscle coordination (Clark et al., 2010) in SKG. In addition, the resulting peak knee flexion varied between subjects as opposed to the consistent reduced knee flexion in SKG. The absence of RF H-reflex modulation and elevated RF H-reflex suggest the exaggerated monosynaptic quadriceps reflex is a prominent factor for reduced knee flexion in SKG.

In conclusion, our results suggest that quadriceps hyperreflexia is a key factor contributing to diminished knee flexion during swing phase in people with post-stroke SKG. Such information could provide a more accurate and quantifiable factor to evaluate the influence of spasticity in post-stroke SKG. Further work is needed to delineate neural mechanisms of this interaction. The implications of this work suggest that interventions targeted at quadriceps hyperreflexia may improve knee flexion in post-stroke SKG.

---

**Table S1: Demographic information of participants with post stroke stiff-knee gait including age, gender, effected side, prescribed medication, ankle-foot orthosis usage, walking speed on the treadmill and average peak knee flexion during walking.**

